# Synthetic genome defenses against selfish DNA elements stabilize engineered bacteria against evolutionary failure

**DOI:** 10.1101/419283

**Authors:** Peng Geng, Sean P. Leonard, Dennis M. Mishler, Jeffrey E. Barrick

## Abstract

Mobile genetic elements drive evolution by disrupting genes and rearranging genomes. Eukaryotes have evolved epigenetic mechanisms, including DNA methylation and RNA interference, that silence mobile elements and thereby preserve the integrity of their genomes. We created an artificial reprogrammable epigenetic system based on CRISPR interference to give engineered bacteria a similar line of defense against transposons and other selfish elements in their genomes. We demonstrate that this CRISPR interference against mobile elements (CRISPRi-ME) approach can be used to simultaneously repress two different transposon families in *Escherichia coli*, thereby increasing the evolutionary stability of costly protein expression. We further show that silencing a transposon in *Acinetobacter baylyi* ADP1 reduces mutation rates by a factor of five, nearly as much as deleting all copies of this element from its genome. By deploying CRISPRi-ME on a broad-host-range vector we have created a generalizable platform for stabilizing the genomes of engineered bacterial cells for applications in metabolic engineering and synthetic biology.

**Significance:** Engineered cells often cease to function or lose productivity when mutations arise in their genomes. Mobile DNA elements, such as transposons, are major sources of these inactivating mutations. Eukaryotic genomes have evolved flexible epigenetic defenses against mobile DNA that help them to maintain genome integrity, but bacteria do not possess comparable silencing systems. We developed a synthetic control system based on CRISPR interference that can be used to give bacterial cells a reprogrammable line of defense against selfish DNA elements in their genomes. We show that this system effectively represses multicopy transposons and multiple families of transposons. Limiting selfish DNA proliferation within a genome in this way improves the reliability of genetically engineered functions in replicating bacterial cell populations.

## Introduction

Unwanted evolution is a foundational challenge for the many areas of biotechnology that rely on genetically engineered organisms (1–3). Engineered cells are often less fit than their wild-type progenitors because they divert resources away from cellular replication or otherwise perturb normal physiological processes (4). Mutations will spontaneously arise in the genomes of some cells in a population that disrupt a DNA-encoded function. Cells with these ‘failure mutations’ often have a significant competitive growth advantage because the engineered burden has been lifted. They will out-replicate the original engineered cells, resulting in a progressive reduction in the performance of the cell population over time (5–8). When the aim is to maximize the production of a recombinant protein or a chemical product, these evolutionary failure modes will reduce yields and limit the useful lifetimes of engineered cells. Genetic instability due to evolution is particularly a problem if there is a large fitness burden for the engineered function during the many cell divisions needed during scale-up of a process to an industrial bioreactor (9).

Simple transposons known as insertion sequence (IS) elements are the dominant source of failure mutations in many engineered bacterial cells (5–8). IS elements are minimal selfish DNA elements: they may consist of just a single transposase gene flanked by inverted repeats (10). They can cause mutations directly, when new IS copies insert into a target DNA site by cut-and-paste or copy-and-paste mechanisms, and indirectly, when recombination between multiple copies of the same IS element leads to deletions or genome rearrangements. Nicking or cleavage of the chromosome by transposases may also induce mutagenic DNA damage responses (11), and transposase binding to the β sliding clamp of the DNA polymerase holoenzyme (12) also has the potential to decrease the fidelity of DNA replication. IS elements can rapidly proliferate within genomes, and they can invade new cells when they are incorporated into DNA elements that mediate horizontal gene transfer, such as conjugative plasmids. Thus, many bacterial genomes harbor multiple copies of several different IS families (13, 14).

If one could silence gene expression from all IS elements that are present in a bacterial host, it would be expected to significantly improve the stability of an engineered function in that cell. Yeast and other eukaryotes have evolved a wide array of epigenetic mechanisms—including DNA methylation, chromatin remodeling, and RNAi—that protect the integrity of their genomes (15). These pathways operate in a flexible manner that enables them to simultaneously silence diverse families of established selfish elements and adapt to newly arrived elements. Many bacteria have defenses, such as RNA-guided nucleases (e.g., CRISPR-Cas9) and restriction-modification systems, that protect their genomes from invasions of new phages, plasmids, and mobile genetic elements (16). However, bacteria do not have a general capacity to silence selfish elements after they have become entrenched in their genomes that is akin to what takes place in eukaryotes.

Perhaps due to this limitation, many IS elements in bacteria have evolved regulatory mechanisms that repress their own activity (17, 18). Presumably, this self-limiting strategy evolved to prevent an IS element from endangering its own survival by overly proliferating within the host genome and causing deleterious mutations in an entire cell population. For example, an antisense RNA is transcribed from the IS*10* transposase reading frame that binds to and inhibits translation of a transposase mRNA produced by the same or any other copy of IS*10* in the genome. Host factors also impact transposition (17, 18). For instance, *dam* methylation inhibits IS*10* transposition, and the nucleoid-like protein IHF facilitates IS*10* transposition. These interactions often modulate transposition so that it is restricted to certain times during DNA replication or to when cells experience stress. These regulatory interactions may represent ‘domestication’ of an IS element such that there is indirect selection to maintain its activity in a bacterial genome because it increases the rates of certain types of beneficial mutations (19, 20), including those that disrupt burdensome plasmids and transgenes added to the genome by human engineering.

Here we describe a reprogrammable plasmid system that prevents evolutionary failures caused by transposons and other mobile elements in genetically engineered bacterial genomes. Our system takes advantage of recently developed CRISPR interference methods (21) and a broad-host-range plasmid backbone (22) that both function in a wide range of bacterial species. Because CRISPR interference can be programmed to bind to and repress specific DNA sequences, one can silence all copies of an active transposon or other selfish element family in a bacterial host genome by adding a single guide RNA to the plasmid. We show that our system for targeting CRISPR interference against mobile elements (CRISPRi-ME) effectively silences multicopy IS elements and multiple families of IS elements. CRISPRi-ME increased the lifetime of burdensome protein expression in *Escherichia coli* and reduced mutation rates in A*cinetobacter baylyi* ADP1. The CRISPRi-ME system can be used to give diverse bacterial species a new system of epigenetic protection against pervasive genomic parasites that cause mutations, thereby improving the reliability of genetically engineered versions of these cells.

## Results

### CRISPR interference from a broad-host range vector

The CRISPRi-ME system uses one or more small guide RNAs (sgRNAs) to target a catalytically dead Cas9 nuclease (dCas9) to bind to specific DNA sequences that silence mobile elements (**Fig. 1**). To implement CRISPRi-ME we started with a well-characterized CRISPR interference (CRISPRi) system that we had previously ported onto a broad-host-range plasmid vector (22). The basic CRISPRi design is derived from plasmids pTargetF and pCas, which were designed for efficient multiplex genome editing in Enterobacteria (23). We substituted a dCas9 gene with the canonical deactivating mutations into this system (21, 24). Then, we added these components to a plasmid backbone derived from the broad-host range expression vector pMMB67EH (25). It contains a low- to medium-copy-number (10-20 plasmids per cell) RSF1010-derived origin of replication that has been shown to function robustly in diverse bacterial species (26), including *E. coli* (27), *Pseudomonas aeruginosa* (28), *Acinetobacter baumanni* (29), and *Snodgrassella alvi* (22). The pMMB67EH backbone also contains an origin of transfer (*oriT*) that enables this plasmid to be conjugated into diverse bacteria.

**Figure. 1.**
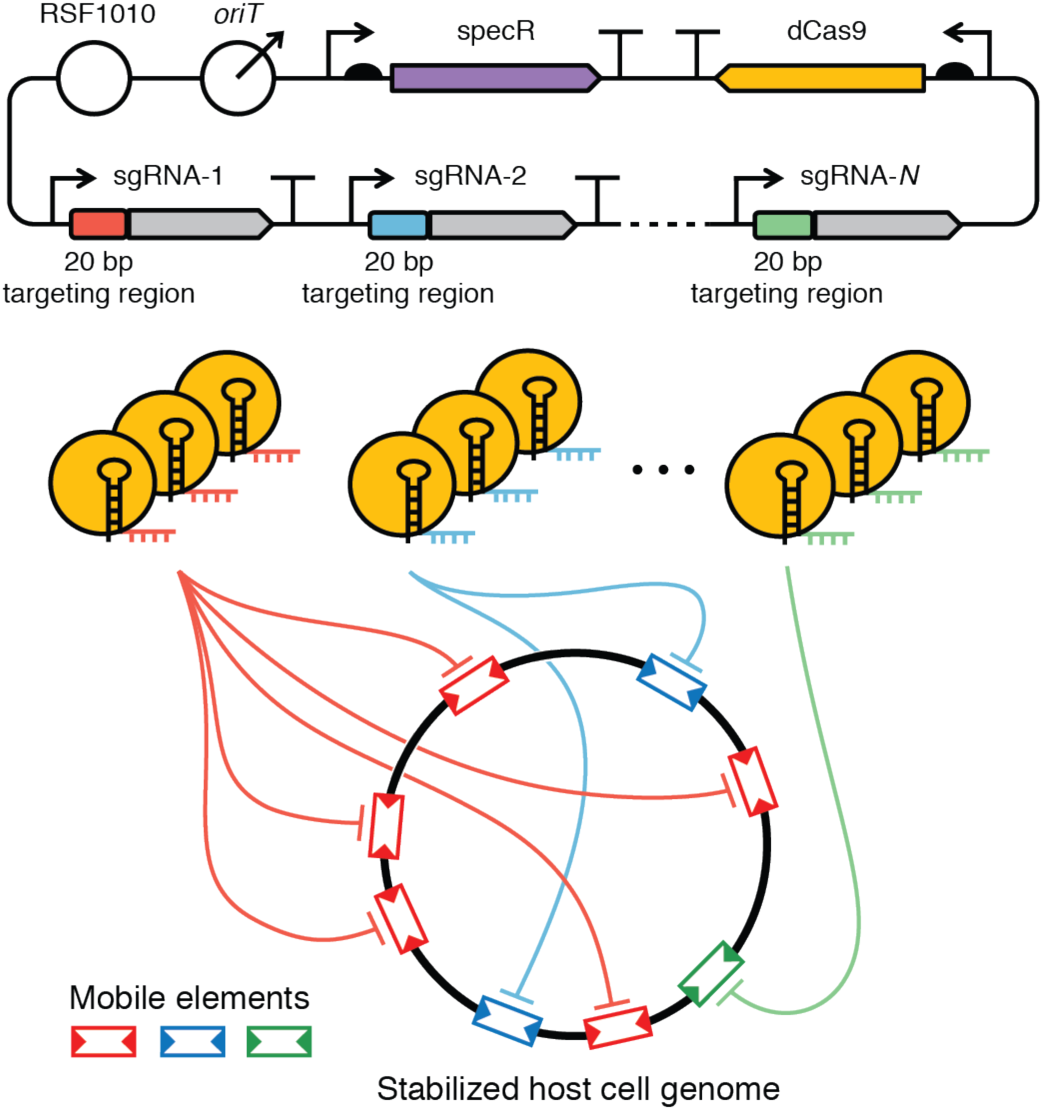
CRISPRi-ME stabilization of a genome against mobile element instability. In the CRISPR interference with mobile elements (CRISPRi-ME) system, one expresses the catalytically inactive dCas9 protein and one or more small guide RNAs (sgRNAs) targeting it to repress genes (e.g., transposases) required for the mobilization of different selfish element families in a host cell. Repressing the activity of mobile DNA elements prevents mutations that commonly inactivate genes required for an engineered function. The pictured configuration uses a broad-host-range plasmid based on the RSF1010 replicon that functions in diverse bacterial species. Plasmid maps in this and other figures are represented using SBOL visual glyphs (69).

To create a customized CRISPRi-ME system (**Fig. 2**), one starts with a plasmid with a single sgRNA targeting unit, a plasmid with the dCas9 transcriptional unit, and a plasmid with the pMMB67EH backbone. Different promoters may be needed to drive transcription of the sgRNA and dCas9 genes to achieve optimal function in different bacterial species, as described in the following sections. Assembly of a CRISPRi-ME plasmid proceeds by first creating one or more variants of the sgRNA plasmid for each targeted mobile element (e.g., IS element transposase) (**Step 1**) and joining them together into a multiple sgRNA targeting cassette plasmid when multiple mobile element families are to be targeted from one CRISPRi-ME plasmid (**Step 2**). Then, all sgRNAs, the dCas9 transcriptional unit, and the pMMB67EH backbone are combined in a final assembly step (**Step 3**). The final CRISPRi-ME plasmid can be purified and transformed into the bacterium of interest or directly transferred into a recipient cell from an *E. coli* strain that encodes the required conjugation machinery (e.g., MFDpir) (30).

**Figure. 2.**
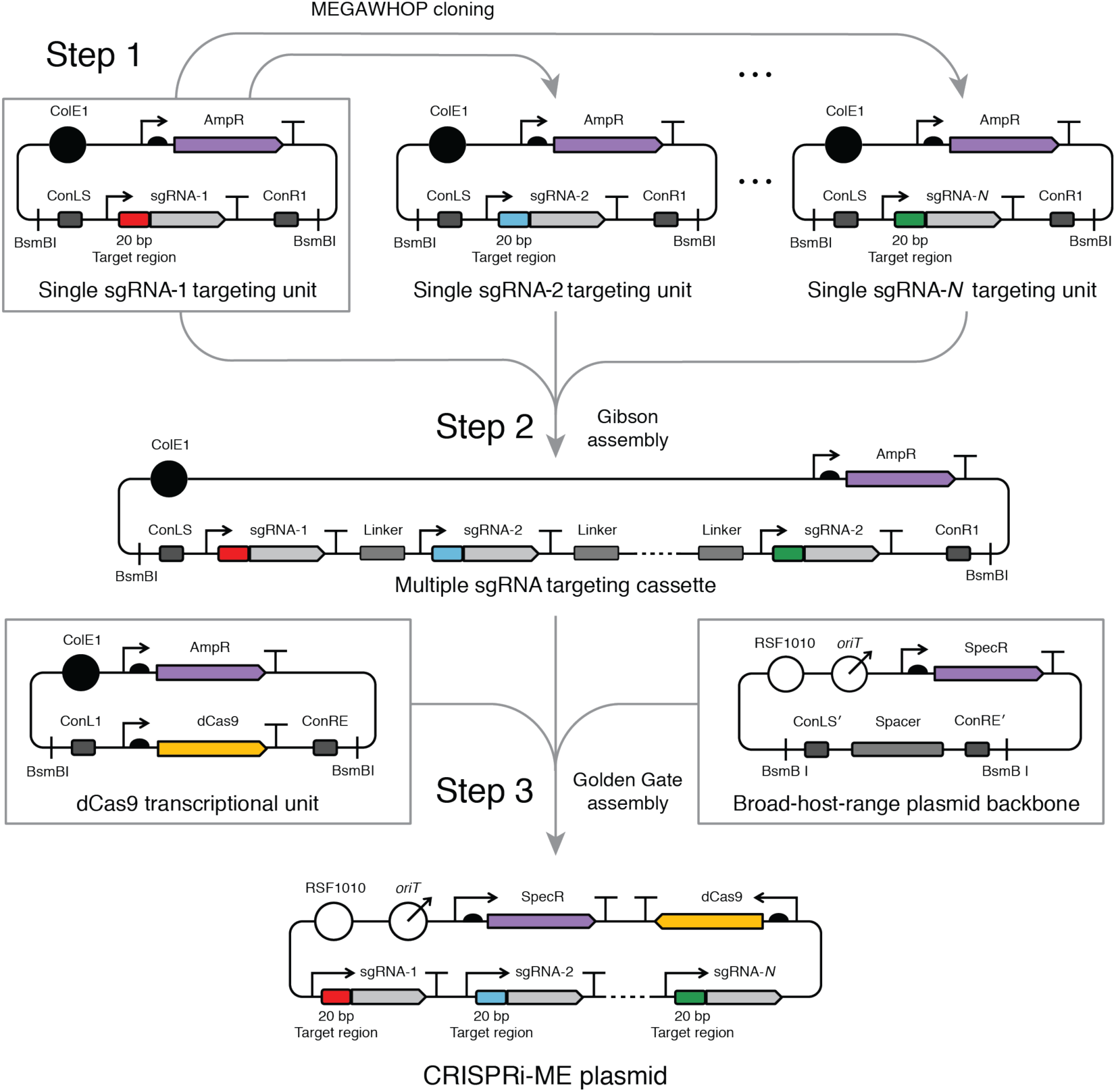
Construction of CRISPRi-ME transcriptional units and plasmids. To create a broad-host-range CRISPRi-ME plasmid, one first constructs a series of plasmids containing individual sgRNA transcriptional units targeted to different mobile elements by changing the 20-base-pair target region of a template plasmid by a method such as MEGAWHOP cloning (**Step 1**). Next, the sgRNA transcriptional units from each of these plasmids are composed into a multiple sgRNA targeting cassette by a sequence-independent cloning method such as Gibson assembly through the addition of unique linker sequences between sgRNA units (**Step 2**). Finally, the dCas9 transcriptional unit and the multiple sgRNA targeting cassette are assembled onto the broad-host-range (RSF1010) plasmid backbone by BsmBI Golden Gate Assembly (**Step 3**). The three boxed plasmids are provided as genetic parts for implementing a custom CRISPRi-ME system. Multiple versions of the single sgRNA targeting unit and dCas9 transcriptional unit plasmids, with different promoters driving sgRNA and dCas9 expression, were created and tested to achieve optimal function in two different bacterial species in this study.

### CRISPRi-ME stabilizes burdensome protein expression in *E. coli*

We first tested how effective our CRISPRi system was at silencing gene expression in *E. coli.* We used previously validated promoters to drive expression of each CRISPRi component (21–23, 31): the native *Streptococcus pyogenes* Cas9 promoter for dCas9 and the constitutive synthetic promoter pJ23119 for each sgRNA (**Fig. 3A**). To test the function of this CRISPRi system on the pMMB67EH plasmid backbone, superfolder GFP (sfGFP) under the control of the native *glpT* promoter was integrated into the genome of a reporter strain of *E. coli*. This CRISPRi configuration strongly repressed expression of sfGFP when an sgRNA targeting this gene was used (>90%) whereas there was no repression with an off-target sgRNA (**Fig. 3B**).

**Figure. 3.**
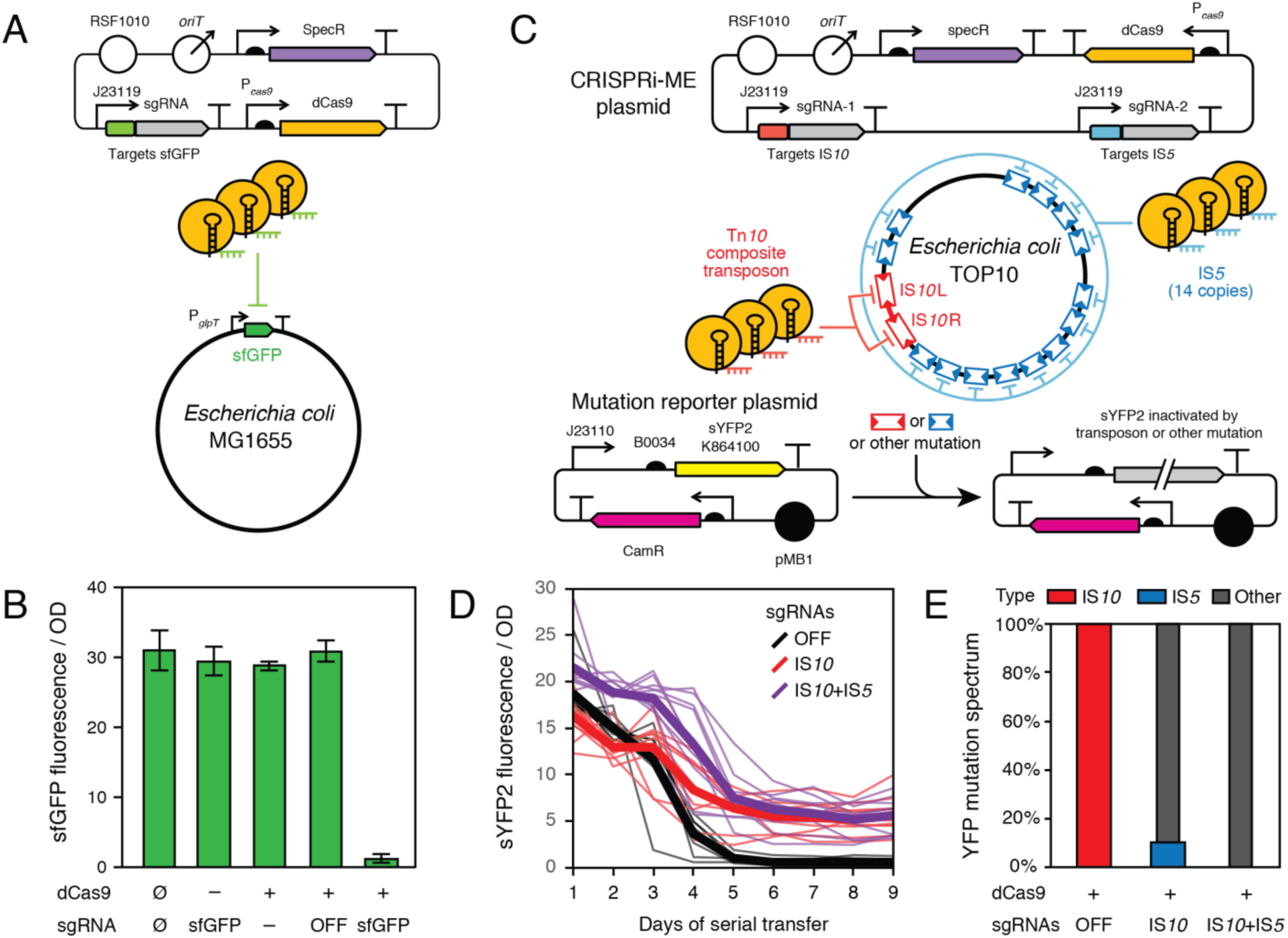
CRISPRi-ME prevents evolutionary failure of a burdensome plasmid in *E. coli*. (A) Design of experiment testing CRISPRi-mediated silencing of a genomically encoded sfGFP reporter gene in *E. coli* MG1655 from a broad-host-range plasmid with an RSF1010 origin. (B) Expression of sfGFP is repressed by this CRISPRi configuration when the sgRNA is targeted to this gene versus in cells with no plasmid (Ø) and in control experiments with plasmids missing either dCas9 or the sgRNA or with dCas9 and an off-target sgRNA (OFF). Wild-type *E. coli* exhibits no fluorescence. Error bars are standard deviations from nine biological replicates. (C) Design of experiment targeting CRISPRi-ME against multicopy IS*10* and IS*5* transposons in the *E. coli* TOP10 genome to extend the evolutionary lifetime of burdensome sYFP2 expression from a high-copy plasmid. (D) sYFP2 fluorescence was monitored over multiple days of serial transfer and regrowth in ten independent cell populations with each CRISPRi-ME plasmid (thin lines). The mean for each treatment at each time point is also shown (thick lines). The CRISPRi-ME plasmids tested contained an off-target sgRNA (OFF), an sgRNA targeting IS*10*, or two sgRNAs targeting IS*10* and IS*5*. (E) Types of mutations that led to a loss of sYFP2 fluorescence in cells containing each CRISPRi-ME plasmid. One evolved sYFP2 plasmid per population was isolated and analyzed at the conclusion of the experiment shown in D.

To determine if CRISPRi-ME could prevent a mobile element from causing inactivating mutations in an engineered DNA sequence, we added this synthetic genetic control system to an *E. coli* TOP10 strain containing plasmid pSB1C3-sYFP2 (**Fig. 3C**). This is a high copy number plasmid constructed from BioBrick parts (32) that strongly expresses a super yellow fluorescent protein variant (sYFP2) (33). In preliminary experiments with this strain, we found that mutant cells with IS*10* element insertions that inactivated sYFP2 expression rapidly arose and outcompeted fluorescent cells. IS*10* is found in two copies that flank the Tn*10* composite transposon in the genome of the host *E. coli* strain (34), and it is known to have strong specificity for certain target site sequences (35). In agreement with this expectation, every independently derived non-fluorescent mutant had an IS*10* insertion at precisely the same site early in the sYFP2 reading frame. We found that editing this target sequence could prevent IS*10* from inserting at this site. When using this edited plasmid, mutations that eliminated the burden of sfYFP expression still arose, but now they were either point mutations or insertions of an IS*5* element. IS*5* is found in 14 copies in the TOP10 genome. It preferentially inserts at sites matching the four-base sequence YTAR (36), which occur many times throughout the engineered DNA sequence, and inactivating IS*5* element insertions were found to occur at various different positions in the construct.

From these preliminary results, we expected that adding a sgRNA targeting the transposase of IS*10* would eliminate the dominant failure mode of pSB1C3-sYFP2 and that adding another sgRNA targeting IS*5* might further stabilize the function of this engineered plasmid. We compared the evolutionary stability of fluorescence in *E. coli* TOP10 cells containing the pSB1C3-sYFP2 plasmid and either a CRISPRi-ME off-target plasmid control, a CRISPRi-ME anti-IS*10* plasmid, or a CRISPRi-ME anti-IS*10*+anti-IS*5* plasmid (**Fig. 3D**). The growth process for each replicate cell population started from culturing an entire single colony that was brightly fluorescent. All cells from the colony were transferred and grown in liquid cultures overnight to saturation (∼35 cell doublings). Then each population was diluted 1:1000 into fresh medium and allowed to regrow for 24 hours (∼10 additional cell doublings per day). Most of the original fluorescence was lost in populations of the CRISPRi-ME off-target strain by the fourth day (∼65 cell doublings), and the fluorescence was totally lost after the sixth day (∼85 cell doublings). In the CRISPRi transposon repressed strains, the fluorescence expression was more stable. Especially in the IS*10*+IS*5* dual repression strain, the host cells were still expressing at least half level the original fluorescence after the fourth day (∼65 cell doublings).

We isolated plasmids from non-fluorescence cells at the end of experiments and sequenced them to determine what types of mutations were responsible for inactivating sYFP2 expression in each case (**Fig. 3E**). In the wild-type *E. coli* strain, sYFP2 was inactivated in 10/10 cases by IS*10* insertions. In contrast, no IS*10* insertions were found in any of the strains with CRISPRi-ME systems containing an anti-IS*10* sgRNA. An IS*5* insertion was found in 1/10 nonfluorescent plasmids from the CRISPRi-ME strain with only the anti-IS*10* sgRNA. In the dual anti-IS*10* and anti-IS*5* sgRNA strain, no IS element insertions in the sYFP2 gene were found. Therefore, by silencing multiple copies of the same IS family and multiple IS families, the CRISPRi-ME system essentially eliminated evolutionary failures due to selfish DNA elements for this construct.

### CRISPRi-ME reduces mutation rates in *A. baylyi* ADP1

The RSF1010 plasmid origin and dCas9 repression system should enable the CRISPRi-ME system to repress mobile elements in a wide variety of bacterial species. To demonstrate its effectiveness in another context we adapted CRISPRi-ME for use in *Acinetobacter baylyi* ADP1. This γ- proteobacterium is of interest in biotechnology due to its natural transformability and metabolic versatility (37–39). Acinetobacter species are more closely related to pseudomonads than they are to enterobacteria (40), and typical *E. coli* plasmids with ColE1-type origins do not replicate reliably in *A. baylyi* (41, 42). The plasmid pMMB67EH, which is the source of the RSF1010 replicon employed in CRISPRi-ME, has been shown to replicate in the related species *Acinetobacter baumanii* (43). Many Acinetobacter species have native type I CRISPR systems in their genomes (44), but neither genome editing nor control of gene expression with Cas9-based systems has been demonstrated previously in this genus to our knowledge.

We found that the RSF1010 backbone used in CRISPRi-ME reliably replicated in *A. baylyi* ADP1. However, it was necessary to change the promoters driving expression of dCas9 and sgRNAs in order for CRISPRi to function effectively from this platform in ADP1 (**Fig. 4A**). For dCas9 we used a promoter that has been used to drive the *tdk* gene, which is used as a counter-selectable marker in this organism (45). For the sgRNA, we used the T5 promoter, which has previously been shown to yield robust constitutive expression in ADP1 (46). With these modifications, there was near-complete repression of an sfGFP gene integrated into the chromosome when the CRISPRi-ME plasmid was used with an on-target sgRNA (**Fig. 4B**). *A. baylyi* ADP1 has six copies of one type of transposable element, IS*1236*, and this element is a dominant source of genetic instability in this strain (47–49). Therefore, we designed a CRISPRi-ME plasmid that represses the IS*1236* transposase (**Fig. 4C**). To determine if silencing IS*1236* stabilized the ADP1 genome against evolution, we used Luria-Delbrück fluctuation assays to measure mutation rates. Loss-of-function mutations in a copy of the *tdk* counterselectable marker inserted into the bacterial chromosome confer resistance to the chain-terminating base analogue, azidothymidine (AZT) (39). Therefore, the mutation rate to AZT resistance yields an aggregate estimate of the risk that an engineered DNA construct inserted into the ADP1 genome has of becoming inactivated by IS*1236* activity or by other mutations.

**Figure. 4.**
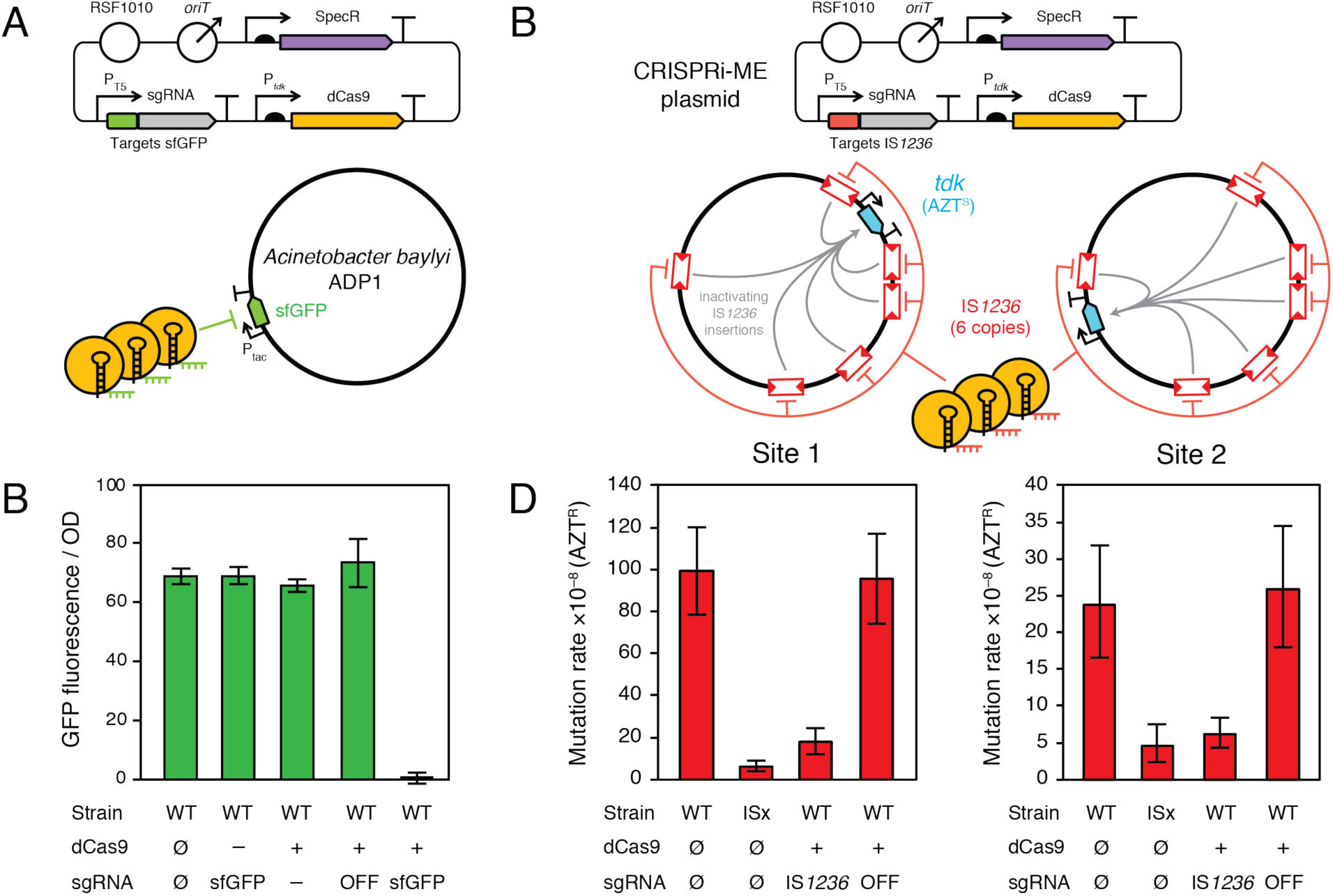
CRISPRi-ME reduces the rates of inactivating mutations in *A. baylyi*. (A) Design of experiment testing CRISPRi-mediated silencing of genomically encoded sfGFP in *A. baylyi* ADP1 from a broad-host-range plasmid with an RSF1010 origin. (B) Expression of sfGFP is repressed by this CRISPRi configuration when the sgRNA is targeted to this gene versus in cells with no plasmid (Ø) and in control experiments with plasmids missing either dCas9 or the sgRNA or with dCas9 and an off-target sgRNA (OFF). Error bars are standard deviations from nine biological replicates. (C) Design of experiment using CRISPRi-ME to silence IS*1236* in *A. baylyi* ADP1. A counter-selectable *tdk* mutational reporter gene was integrated into the ADP1 chromosome at different sites in two strains. Expression of the *tdk* gene results in toxic incorporation of AZT during DNA replication. If an inactivating mutation occurs in *tdk*, it enables cells to grow on selective agar containing AZT. (D) Mutation rates to AZT resistance in strains containing anti-IS*1236* and off-target CRISPRi-ME plasmids estimated from Luria-Delbrück fluctuation tests. ISx is a variant of wild-type *A. baylyi* ADP1 (WT) with all five IS*1236* elements deleted from its chromosome. Error bars are 95% confidence intervals.

We measured mutation rates to AZT resistance in two ADP1 host strains that had the *tdk* mutational reporter gene integrated at different locations in the bacterial chromosome (**Fig. 4D**). We found that the presence of the anti-*IS1236* CRISPRi-ME system reduced the rates of inactivating mutations in the *tdk* gene by a factor of five at both sites whereas strains with the off-target sgRNA exhibited no change in mutation rates. The five-fold reduction in mutation rates indicates that there is near-complete suppression of IS*1236* activity, as mutation rates in strains with the CRISPRi-ME system were almost as reduced as they were in a positive-control ‘clean-genome’ ADP1-ISx strain in which all six IS*1236* elements were deleted from the genome (49).

## Discussion

In this study, we developed and employed CRISPR interference against mobile elements in bacteria. This CRISPRi-ME approach reduced the detrimental effects of IS elements on the continued production of target biomolecules and significantly stabilized genetically engineered DNA sequences. Specifically, we prevented inactivating mutations that result in the loss of burdensome protein expression from a plasmid in the *E. coli*, and we reduced mutation rates in the bacterial chromosome by as much as 5-fold in *A. baylyi*. Because CRISPRi-ME employs broad-host-range components (the RSF1010 replicon and the dCas9 catalytically inactivated RNA-guided nuclease), it can be readily reprogrammed to function in diverse bacterial species.

To completely prevent loss-of-function mutations generated by insertion sequences, ‘clean-genome’ bacterial strains have been constructed in which one or more IS element families and sometimes other selfish elements, like prophage, have been deleted from the chromosome. Examples of clean-genome strains include *Escherichia coli* MDS42 (5), *Pseudomonas putida* EM383 (50), *Corynebacterium glutamicum* WJ004 and WJ008 (51), and *Acinetobacter baylyi* ADP1-ISx (49). Engineering projects that begin in these strain backgrounds do not have to worry about IS element activity. However, many strains of bacteria used in research and industrial applications already exist that have been subjected to extensive genome editing efforts or directed evolution during which many beneficial mutations have accumulated in their genomes (52–55). Preventing IS elements from compromising the functions of these highly engineered strains is nontrivial. One must either identify the mutations that are important for the strain’s function and re-engineer them into a clean-genome strain background or repeat the process of sequentially deleting selfish elements from the engineered strain’s genome, which is labor-intensive (5, 49).

In eukaryotic cells that have efficient nonhomologous end joining (NHEJ), it is possible to simultaneously inactivate many members of a single selfish DNA element that contributes to genome instability by targeting them for cleavage with an RNA-guided nuclease (56). NHEJ processes do not exist or are inefficient in most bacteria (57), including *E. coli* (58), but it may be possible in the future to heterologously express a NHEJ system to achieve multiplex editing that could be used to inactivate selfish DNA elements in bacterial genomes (59). Alternatively, the process of re-cleaning a new bacterial genome can be accelerated by using a related clean-genome strain as a donor for transduction and existing multiplex genome editing methods (60).

The CRISPRi-ME approach is to silence the expression of mobile elements, rather than to delete them from the bacterial chromosome. It resembles how eukaryotic genomes have evolved defenses to maintain genome integrity against abundant selfish DNA elements in their genomes. In the context of bacterial genetic engineering, the CRISPRi-ME system can be used to rapidly prototype whether silencing a particular mobile element family will increase the stability of an engineered function before investing in the time-consuming process of deleting all of its copies from a genome. We did not observe a growth rate cost for adding this exogenous silencing control system to cells, so it could also be directly useful for stabilizing certain bioproduction processes. Recently Tn*7* and ICE element based tools for integrating CRISPRi systems into bacterial genomes have become available (61). CRISPR-iME could be implemented using these systems when maintaining a genetic control plasmid is not desirable and for adding compatibility with even more bacterial species. CRISPRi-ME gives bacteria a synthetic line of defense against endogenous mobile DNA elements, thereby stabilizing the function of genetically engineered cells.

## Materials and Methods

### Bacterial strains and growth conditions

*E. coli* strains were cultured at 37°C in Lysogeny Broth (LB) (10 g NaCl, 10 g tryptone, and 5 g yeast extract per liter). We used *E. coli* DH5α for all cloning steps. *A. baylyi* was cultured in LB at 30°C. Both bacteria were incubated with orbital shaking at 200 r.p.m. over a 1-inch diameter. Media amendments were added at the following concentrations when specified: kanamycin (Kan), 50 µg/ml; spectinomycin (Spec), 60 µg/ml; carbenicillin (Crb), 100 µg/ml; chloramphenicol (Cam), 20 µg/mL; 3′-azido-2′,3′-dideoxythymidine (AZT), 200 µg/ml.

### Broad-host-range CRISPRi platform

Lee et al. constructed a versatile yeast toolkit (YTK) for Golden Gate assembly of plasmids (62), and we extended it to enable genetic engineering of bacteria from the bee gut microbiome (BTK) (22). These kits designate particular restriction enzyme overhangs for promoters, coding sequences, terminators, and connecters that allow plasmids to be hierarchically assembled using Golden Gate assembly. We followed the basic design principles used in the BTK for CRISPRi-ME plasmid construction as illustrated in **Fig. 2**. The five component plasmids needed to assemble the CRISPRi-ME systems validated in *E. coli* and *A. baylyi* in this study have been submitted to the Addgene plasmid repository. Their sequences are provided in **Dataset S1**.

The first two component plasmids for the single sgRNA targeting unit and dCas9 transcriptional unit plasmids were created by cloning these genes into pYTK095, which has a ColE1 origin (62). The sgRNA targeting unit plasmid contains connectors ConLS and ConR1 flanking a sgRNA transcriptional unit. Megaprimer PCR of Whole Plasmids (MEGAWHOP) cloning (63) was used to change the 20-base sgRNA target region in this plasmid to the sequences given in **Table S1** for different experiments. Each *E. coli* sgRNA was checked for potential off-target binding sites in the bacterial genome using the Cas-Designer web tool (64). For the multiple target CRISPRi system, the sgRNA transcriptional units from two such plasmids were assembled into the pYTK095 plasmid backbone using Gibson assembly with an arbitrary DNA linker added between them to maintain terminal ConLS and ConR1 linkers. The dCas9 transcriptional unit is flanked by ConL1 and ConRE connectors in its plasmid. The coding sequence for dCas9 was derived from plasmid pdCas9 (24) with the removal of an internal BsmBI site. The sgRNA and dCas9 transcriptional units were assembled together with the RSF1010 origin from pMMB67EH using BsmBI Golden Gate assembly.

### GFP repression assays

An *E. coli* MG1655 derivative constitutively expressing sfGFP from the chromosome was created using λ Red recombination (65). Briefly, we generated a DNA fragment with the native *E. coli glpT* promoter controlling sfGFP linked to an adjacent chloramphenicol resistance gene via PCR reactions that also added 50-bp extensions homologous to regions adjacent to the *lacZ* gene. This product was electroporated into cells induced to express the λ Red proteins from plasmid pKD46 as previously described (66). A fluorescent colony was selected on LB-Cam agar and then cured of the temperature-sensitive pKD46 plasmid to isolate strain MG1655-sfGFP.

For *A. baylyi* we used natural transformation to add a similar cassette to the chromosome at a neutral location (Site 2) as previously described (49). Briefly, a double-stranded DNA fragment which contained sfGFP under control of the Tac promoter, a chloramphenicol resistance gene, and two 1-kb chromosomal flanking homology regions was constructed by PCR. Then, *A. baylyi* ADP1 was transformed with this DNA fragment as previously described (67). A fluorescent colony was selected after plating these cells on LB-Cam agar and designated strain ADP1-sfGFP.

CRISPRi-ME plasmids were transformed into MG1655-sfGFP and ADP1-sfGFP to test the effectiveness of gene silencing. Entire colonies were scraped from agar plates and inoculated into 10 mL of LB. After incubation for 12 hours, the absorbance at 600 nm (OD600) and fluorescence (excitation 488 nm, emission 525 nm) were measured for 100 μl samples taken from these cultures using a Tecan Infinite M200 PRO microplate reader. The off-target sgRNA used in these tests was targeted to a different fluorescent protein variant, GFP optim-1 (22).

### Monitoring plasmid failure

The *E. coli* mutational reporter plasmid was constructed by BioBrick assembly of promoter (J23100), ribosome binding site (B0034), and sYFP2 fluorescent protein (K864100) parts obtained from the iGEM Registry of Standard Biological Parts (32). There were two six-base-pair repeats (TACTAG) located upstream and downstream of the ribosome binding site in this initial plasmid that mediated a deletion that dominated among the mutations leading to non-fluorescent cells after IS*10* silencing in preliminary experiments. To eliminate this mutational hotspot, the upstream repeat copy was modified to GTATAG to create the reporter plasmid used in our experiment.

For each strain tested in the decay experiment, ten different strongly fluorescent colonies on a LB agar plate were each transferred into test tubes containing 5 ml of LB. After 24 hours of growth (designated day 1 of serial transfer), 5 µl of culture was transferred from each test tube into 5 ml of fresh LB in a new test tube. This procedure was repeated for eight additional days. Fluorescence (excitation 495 nm, emission 530 nm) and OD600 were monitored as described above. The off-target sgRNA control in this experiment was targeted to the *A. baylyi* ADP1 IS*1236* sequence.

### Mutation rate measurements

For each strain, an initial overnight culture was grown in LB-Spec for strains carrying a CRISPRi-ME plasmid or LB for other strains. Then, fourteen independent 100 µl cultures per strain in 18 × 150 mm test tubes in the same media were each inoculated with ∼500 cells from the overnight culture. These new replicate cultures for the fluctuation test were then allowed to grow overnight (∼16 h) to saturation. To estimate the total number of cell numbers in the final cultures, dilutions in sterile saline from two of the tubes were plated on nonselective LB agar plates. The entire volumes of the other twelve tubes were plated separately on selective LB-AZT agar plates. All plates were incubated at 30°C for 24 h, then colony numbers were counted. Mutation rates were estimated from these counts using rSalvador (version 1.7) (68). The off-target sgRNA used in this experiment targeted the GFP optim-1 sequence, as above.

## Acknowledgments

We thank Kelsey Hu and the UT Austin “Microbe Hackers” Freshman Research Initiative stream for their work identifying the evolutionary failure modes of the sYFP2 plasmid and Gabriel Suárez for advice and materials for working with *A. baylyi* ADP1.

## Funding Information

This work was funded by the Defense Advanced Research Projects Administration (HR0011-15-C0095) and the National Science Foundation (CBET-1554179).

**Dataset S1. Sequences of plasmids used to assemble CRISPRi-ME systems and sgRNA target sites in GenBank Flat File Format**

